# Env from EIAV vaccine delicately regulates NLRP3 activation via attenuating NLRP3-NEK7 interaction

**DOI:** 10.1101/2024.11.26.625355

**Authors:** Xing Guo, Cong Liu, Yuhong Wang, Hongxin Li, Saiwen Ma, Lei Na, Huiling Ren, Yuezhi Lin, Xiaojun Wang

## Abstract

The current equine infectious anemia virus (EIAV) vaccine causes attenuation of the inflammatory response to an appropriate level, compared to that produced by virulent EIAV. However, how the EIAV vaccine finely regulates the inflammatory response remains unclear. Using a constructed NLRP3-IL-1β screening system, viral proteins from two EIAV strains (the attenuated vaccine and its virulent mother strain) were examined separately. Firstly, EIAV-Env was screened to direct binding P2X7(R) with notable K^+^ efflux trans-cellularly. Secondly, EIAV-Env was found to bind NLRP3 and/or NEK7 to trigger aggregation of NLRP3-NEK7 to form NLRP3-NEK7 complex in cells. Comparison of the two strains, we observed a significant reduction on vaccine-Env-initiated NLRP3-NEK7 complex formation, with no difference in Env triggering P2X7(R)-mediated ion fluxes. Thirdly, reciprocally mutation on four stable varied amino acids between two strains produced an anticipated outcome on NLRP3-IL-1β-axis activation. As the attenuated vaccine was shown evolved as a natural quasispecies of the virulent EIAV, its precise and adaptable regulation via spatial proximity-dependent intracellular activation might present a “win-win” virus-host adaption, offering an alternative strategy on HIV vaccine development.

**Author Summary:** Here, we report that EIAV-Env mediates NLRP3 inflammasome activation through two distinct pathways. The first pathway involves a transcellular mechanism driven by K+ flux, which couples Env-P2X7 interaction. The second pathway entails direct intracellular binding between Env and NLRP3, promoting the assembly of NLRP3-NEK7 and subsequent inflammasome formation. Notably, we observed a marked difference in NLRP3 inflammasome activation between the vaccine and virulent strains, which was reflected in the extent of Env-mediated NLRP3-NEK7 aggregation. This study not only enhances our understanding of lentivirus-host immune interactions but also contributes to the broader discourse on virus evolution and host-induced inflammation.

## Introduction

Equine infectious anemia virus (EIAV) belongs to the lentivirus family, and shares similar virus-host immunity and pathogenesis with the human immunodeficiency virus (HIV-1) [1–4]. Despite extensive efforts, safe and sustainable HIV-1 vaccines have yet to be developed, primarily due to the presence of latent virus reservoirs and rapid viral evolution [5–7]. In contrast, an attenuated vaccine against EIAV, derived from its virulent progenitor strain, has been successfully developed and widely utilized for decades [3, 8]. Notably, natural EIAV quasispecies in equine populations have also been observed to undergo attenuation with time [9–12]. Through systemic sequencing, our “artificially” attenuated EIAV vaccine has been demonstrated to share similar evolutionary traits with the naturally attenuated EIAV [8, 13]. This suggests that the vaccine can be regarded as a “natural” EIAV that has undergone accelerated evolution. Several studies have demonstrated that the virulent EIAV strain elicits more pronounced inflammatory responses and pathological effects than those produced by the attenuated EIAV vaccine [4, 14]. Specifically, there are significant differences observed in the production levels of interleukin-1 beta (IL-1β), a cytokine tightly regulated by the inflammasome, between the two EIAV strains. The NLRP3-IL-1β inflammatory pathway is well-documented in viral pathogenesis, often triggered by various pathogens through P2X7-dependent signaling [15]. A vast number pathogens are able to trigger NLRP3-IL-1β-axis activation via P2X7-dependent signaling [16–19]. However, it’s noteworthy that certain pathogens activate the NLRP3 inflammasome through ATP- and P2X7-independent pathways [20, 21]. Despite these insights, the mechanism by which EIAV activates the inflammasome and subsequent inflammatory pathways remains unclear. Therefore, the aim of the study is to delve into the molecular profiles of inflammasome activation induced by both the attenuated vaccine strain and the virulent mother strain of EIAV. This investigation seeks to uncover the reasons behind the differential regulation of the inflammasome by these two EIAV strains. Our findings may provide a potential target for fine-tuning the regulation of the NLRP3-IL-1β axis, which could have implications for the development of safe and efficient vaccines against human lentiviruses.

## Results

### The attenuated vaccine triggers markedly lower NLRP3-IL-1β-axis activity than the virulent EIAV

The utilization of the semi-quantitative scoring platform, revealed a notable reduction in inflammatory lesions within the vaccine group compared to the virulent group (S1 Fig and Table 1) [22]. This finding highlights the potential effectiveness of the vaccine in attenuating inflammatory reactions and pathological harm associated with EIAV infection. Moreover, previous studies have demonstrated significant differences in the presence of IL-1β following infection with the EIAV vaccine compared to the virulent strain [4, 23]. These findings further strengthened the understanding of the vaccine’s impact on inflammatory responses and disease progression. Therefore, we further compared the molecular regulation of IL-1β associated pathways by the EIAV vaccine strain and the EIAV virulent strain parallelly *in vivo* and *in vitro*. *In vivo* infections resulted in a consistent elevation of IL-1β induced by EIAV vaccine or by virulent strain, with much lower trajectory induced by vaccine (Fig1A). *In vitro* studies in equine macrophages showed a similar difference between the attenuated and virulent EIAV strains (Fig 1B). Interestingly, the transcription level of pro-IL-1β did not significantly differ between the two groups, indicating that the vaccine robustly regulated the conversion of pro-IL-1β to mature IL-1β (Fig1C). The NLR family proteins play a key role in the regulation of the conversion of pro-IL-1β into mature IL-1β, and we therefore compared the expression levels of NLRP1, NLRP3 or NLRC4 mRNA between the two groups using quantitative RT-PCR. We found that NLRP3 mRNA was significantly enhanced in macrophages following infection with either virulent EIAV or the attenuated vaccine, with no significant differences between the two strains. Moreover, the expression of NLRP1 and NLRP4 mRNA was not induced following infection with either strain of EIAV (Fig 1D). We also visualized the formation of ASC speck, a hallmark of inflammasome activation, in macrophages infected with either the attenuated vaccine or with virulent EIAV (Fig 1E). Using inhibitor (MCC950) to inhibit NLRP3-IL-1β signaling, the expression of IL-1β induced by EIAV was remarkably down-regulated with a similar level as negative control (Fig 1F), further confirming that the NLRP3-IL-1β-axis had been activated by both the virulent strain and by the attenuated vaccine.

**Table 1.** Representative images of inflammatory pathological lesions in lung, liver, spleen, kidney and lymph node (•20) from three equines inoculated with EIAV attentuated vaccine or from three equine infected with EIAV virulent strain

| Group | Animal ID | Scoring on inflammatory pathological lesions |  |  |  |  |
| --- | --- | --- | --- | --- | --- | --- |
|  |  | lung | liver | spleen | kidney | lymph node |
| Vaccine strain | Vac-1 | 0 | 0.5 | 0 | 0 | 0 |
|  | Vac-2 | 0 | 1 | 0 | 0 | 1 |
|  | Vac-3 | 0.5 | 0 | 0 | 0.5 | 1 |
| Virulent strain | Vir-1 | 3 | 2.5 | 2.5 | 2 | 2.5 |
|  | Vir-2 | 2.5 | 2 | 2 | 2 | 3 |
|  | Vir-3 | 3 | 1.5 | 3 | 1 | 2 |

Note: Using the semi-quantitative scoring method that we had applied, inflammatory pathological lesions in five organs from three horses that inoculated with attenuated vaccine or from three horses that infected with virulent strain were scored. "0" indicated a normal organ, "1" indicated mild pathological changes, "2" indicated moderate pathological changes and "3" indicated severe pathological changes. 0.5 score was applied for more delicate comparison in the present study with agreement of two professional pathologists

**Fig 1.**
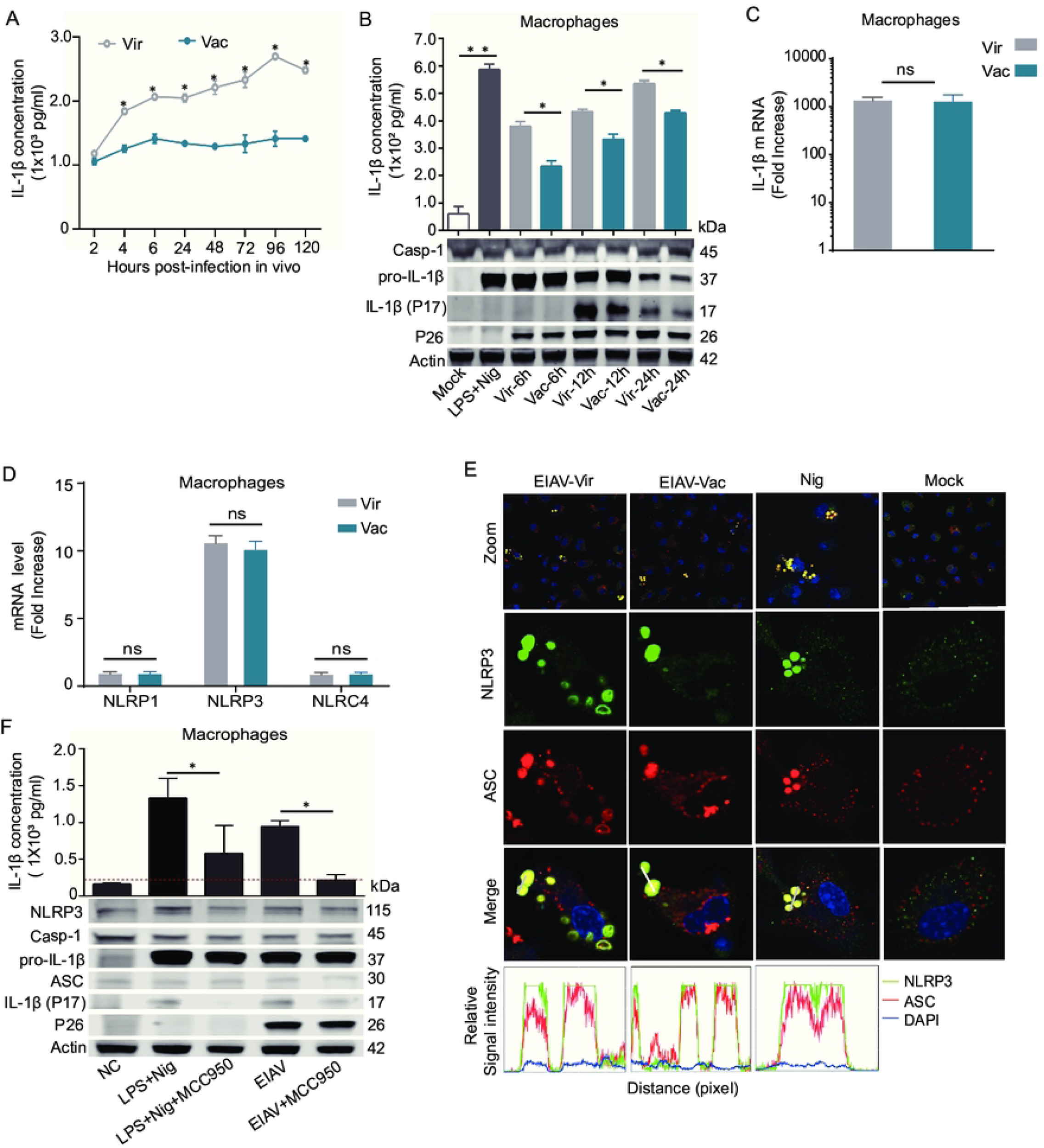
The vaccine EIAV induces markedly lower levels of IL-1β than the virulent EIAV *in vivo* and *in vitro.* **(A)** Quantification of IL-1β levels in the sera of individuals infected with virulent EIAV and vaccine EIAV using ELISA assay. (**B**) Quantification of IL-1β in the supernatants of virulent and vaccine EIAV-infected macrophages. Western blot analysis addressing expression of pro-IL-1β and IL-1β was performed parallelly. (**C-D**) Quantification of mRNA expression of IL-1β (C) and NLR family genes (*NLRP1/NLRP3/NLRC4*) (D) in macrophage cells infected with virulent or vaccine EIAV strains using real-time RT-PCR. (**E**) Visualization of endogenous NLRP3 (green)-ASC (red) puncta formation in macrophages infected with virulent or vaccine strain (12 h *p.i.*) using confocal microscopy (Nigericin-treated macrophages served as the positive control). (**F**) Representative images and statistical analyses from Western blot and ELISA demonstrate the expression levels of IL-1β in equine macrophages treated with MCC950 and EIAV (2 x 10^5^ TCID50, 24 h) or with LPS (10 mg/mL, 6 h) and Nigericin (10 μM, 2 h) as control.

### Screening for EIAV-Env component able to activate the NLRP3-IL-1β-axis interactively

We observed that the activation of the NLPR3 inflammasome and the levels of IL-1β appeared significantly difference between the two EIAV strains. We therefore wanted to screen for any component of EIAV that interacted with any molecule within the NLRP3-IL-1β-axis. We constructed an equine NLRP3-IL-1β-axis screening system (Fig 2A), and found that the viral Env protein was the dominant component of EIAV that induced IL-1β secretion with a dosage-dependent effect in this system (Fig 2B). Furthermore, we confirmed the subunit of Env (Gp90) that activated the NLRP3 inflammasome (Fig 2C). We then visualized the ASC speck formation in *env*- or *gp90*-transfected 293T cells with a confocal microscope to further validate these findings (Fig 2D). Then equine macrophages were infected with two Env-pseudotyped viruses. We observed that secretion of mature IL-1β was induced by both of the Env-pseudotyped viruses, confirming that Env could mediate NLRP3-IL-1β-axis activation (Fig 2E). Next, we sought to determine if vaccine-Env and virulent-Env differed in NLRP3-IL-1β-axis activation. 293T cells were co-transfected with Envs and plasmids of NLRP3-IL-1β screening system. We observed significantly lower IL-1β production in 293T cells transfected with vaccine-*env*, compared with that in 293T cells transfected with virulent-*env* (Fig 2F). From these analyses, we speculated that Env was the main factor affecting the effects of the NLRP3-IL-1β-axis activation induced by EIAV infection.

**Fig 2.**
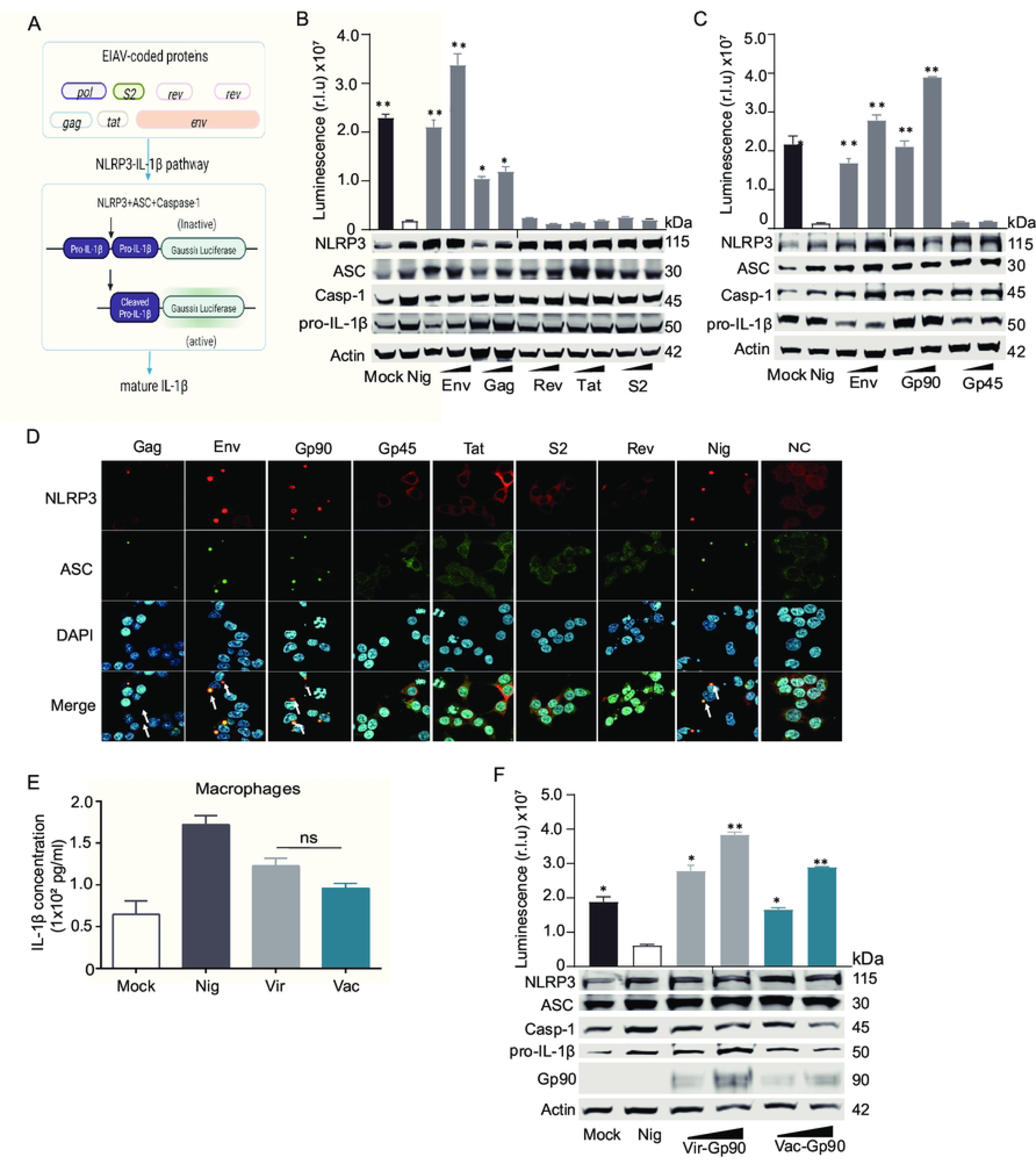
Screening for EIAV-coded proteins involved in the NLRP3-IL-1β-axis activation. (**A**) Schematic of the bioactivity of the NLRP3-IL-1β signaling in 293T cells co-transfected with EIAV-coded proteins *via* a luciferase reporter system. (**B**) 293T cells were co-transfected with the indicated reporter constructs in the presence or absence of EIAV expression plasmids separately. Luciferase activity and reporter construct expression were analyzed as in (**A**). (**C**) Cells were subjected to the same assay protocol as (**B**) but were transfected with *gp90* and *gp45* separately. Analysis was performed as described in (**A**). (**D)** Con-focal microscopy visualization of the formation of NLRP3-ASC puncta (yellow, indicated by white arrows) in 293T cells co-transfected with one of the five EIAV-coded proteins (Env/Gag/S2/Rev/Tat) or with one of the two Env glycoprotein components (Gp90 and Gp45). (**E**) Statistical analysis of ELISA data was conducted to evaluate the expression levels of IL-1β in macrophages infected separately with Env-pseudotyped viruses derived from virulent or vaccine EIAV strains at 24 hours post-infection. F Same as (B) but co-transfected with virulent-*env* and vaccine-*env* separately. (\**P* < 0.05, ** *P* < 0.01, compared to the mock-transfected group).

### Screening target cellular proteins involved in the activation of NLRP3-IL-1β-axis by EIAV-Env

To screen for target cellular protein (s) that are associated with NLRP3 inflammation activation by EIAV-Env, we performed an Env IP-mass spectrometry (IP-MS) (Fig 3A). Three target inflammasome-associated proteins, P2X7(R), NEK7 and NLRP3, were shown to directly interact with Env in virulent EIAV-infected macrophages (Fig 3B and 3C). We further validated this result using a co-immunoprecipitation (Co-IP) assay. We observed that Env co-precipitated with P2X7(R), NEK7 or NLRP3 in both vaccine-*env* or virulent-*env* transfected 293T cells (Fig 3D-3F). Co-localization of Env and P2X7(R), NEK7 or NLRP3 in both groups could be visualized using confocal microscopy (Fig 3G-3I). Therefore, we speculated that the activation of the NLRP3 inflammasome induced by EIAV infection occurred *via* a mechanism dependent on Env/P2X7(R)/NLRP3/NEK7 binding. Interestingly, no disparity between the abilities of Env to bind these proteins was observed between the virulent and vaccine EIAV strains.

**Fig 3.**
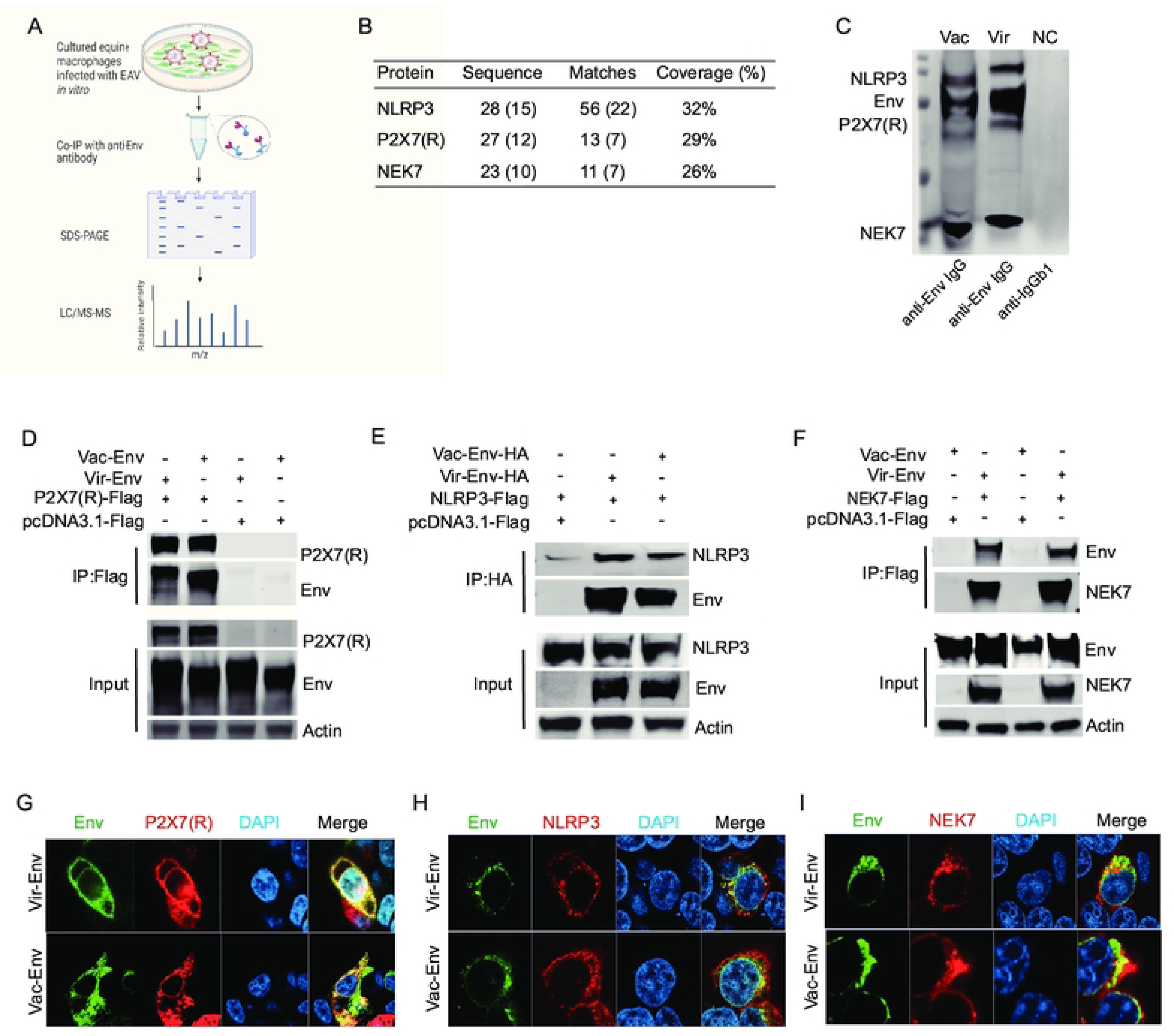
Screening for Env-interacted cellular protein (s) involved in the NLRP3-IL-1β-axis. (**A**) Schematic illustration of the IP-mass spectrometry screening for protein (s) in the NLRP3-IL-1β-axis that interact with EIAV-Env. (**B**) Mass spectrometry analysis of NLRP3, P2X7 (R), and NEK7 peptides following EIAV infection. (**C**) Immunoprecipitation with anti-Env antibodies of endogenous P2X7 (R), NLRP3 or NEK7 in macrophages infected with EIAV stain. (**D**) Co-immunoprecipitation of HA-Env with Flag-NLPR3 in 293T cells transfected with the indicated expression plasmids. (**E**) Same procedure as (D) but cells transfected with Flag-NLRP3 and Flag-*env* (s). (**F**) Same procedure as (D) but cells transfected with Flag-NEK7 and Flag-*env* (s). (**G**-**I**) Co-localization between Flag-P2X7R and HA-Env, Flag-NLRP3 and HA -Env or Flag-NEK7 and HA-Env was examined using confocal microscopy. Scale bars: 10 µM.

### Investigation of the P2X7(R)-regulated K^+^/Ca^2+^ channel status following infection with attenuated vaccine or virulent EIAV

The role of the Env-P2X7(R) interaction in the regulation of the NLRP3 inflammasome was then explored. As an ATP-gated ionic channel [24, 25], the opening of P2X7(R) leads to low intracellular K^+^ concentrations or high intracellular Ca^2+^ concentrations, which then trigger the activation of the NLRP3 inflammasome [17, 24]. We therefore examined whether or not the P2X7(R) ionic channel was activated in EIAV-infected macrophages through comparison of intracellular K^+^/Ca^2+^ concentrations with or without EIAV infection. As excepted, we observed that intracellular K^+^ concentrations rapidly decreased (Fig 4A) in macrophages infected by either the virulent or the vaccine EIAV strain, while the Ca^2+^ concentrations remain stable or slightly increase in time (Fig 4B). Consistent with this result, P2X7(R) knockdown prevented K^+^ efflux in macrophages infected with attenuated vaccine or the virulent EIAV (Fig 4C). To further validate this in 293T cells induced by Env, we pretreated 293T cells using Gly (a K^+^ inhibitor), or grew the cells on medium containing either high K^+^ concentrations and then measured the activation of the NLRP3-IL-1β-axis. We observed that Gly (a K^+^ inhibitor) significantly reduced NLRP3-IL-1β-axis activation in both attenuated Env and virulent Env (Fig 4D), a finding further supported by treatment with high K^+^ medium (Fig 4E). These observations suggest that both virulent-Env and vaccine-Env are able to induce K^+^ efflux-dependent NLRP3-IL-1β-axis activation. Notably, no differences were observed in the Env-mediated K^+^ efflux between the two EIAV strains.

**Fig 4.**
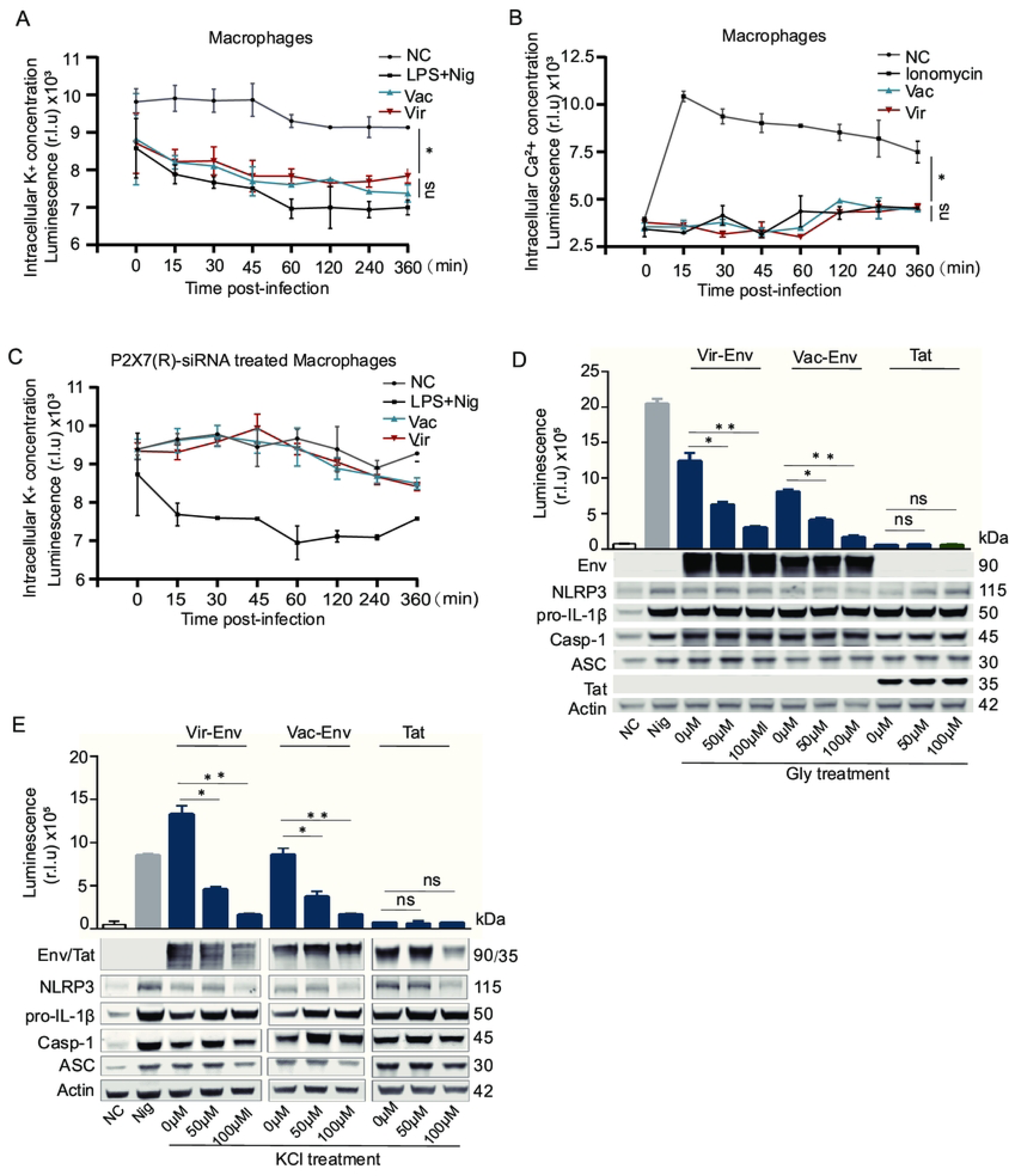
Screening for ion-dependent P2X7(R)-initiated NLRP3-IL-1β-axis activation induced by virulent-Env or by vaccine-Env. (**A**) Time-dependent changes in intracellular K^+^ concentrations in macrophages infected with virulent EIAV or vaccine EIAV. Here NC refers to negative control and LPS+Nig means positive control. (**B**) Time-dependent changes in intracellular Ca^2+^ concentrations in macrophages infected with virulent EIAV or vaccine EIAV. (**C**) Time-dependent changes in intracellular K^+^ concentrations in macrophages pre-treated with P2X7(R) receptor-specific siRNA for 6 h, and then infected with virulent EIAV or vaccine EIAV. (**D**-**E**) Evaluation IL-1β in supernatants of 293T cells co-transfected with virulent-*env* or vaccine-*env* in the presence of increasing doses of the K^+^ efflux inhibitor (Gly, 50 µM,100 µM) (D) and KCl (50 µM,100 µM) (E) (Tat served as a negative control). (\**P* < 0.05, ** *P* < 0.01)

### Vaccine-Env attenuated the NLRP3-NEK7 complex compared with virulent-Env

To explore whether P2X7 (R) contributes to Env-induced NLRP3-IL-1β-axis activation, we generated 293T cell lines stably expressing P2X7(R). The use of P2X7(R)-specific siRNA and a P2X7(R) inhibitor revealed a significant reduction in IL-1β production in cells treated with either the virulent-Env or vaccine-Env, compared to untreated groups. Interestingly, the observed reduction in IL-1β production was similar between cells treated with the attenuated vaccine-Env and those treated with the virulent-Env (Fig 5A and 5B), indicating that the involvement of P2X7(R) in IL-1β regulation is consistent regardless of the virulence of the strain. By elucidating the ability of Env to directly bind NLRP3 or NEK7 in cells, our findings bolster the hypothesis of a P2X7(R)-independent activation mechanism contributing to NLRP3 inflammasome activation during EIAV infection (Fig 3E and 3F). Following the co-transfection of three plasmids (*env*, *NLPR3*, and *NEK7*) simultaneously into 293T cells, a notable increase in NLPR3-NEK7 precipitation was observed in cells transfected with either vaccine-*env* or virulent-*env* compared to the mock-transfected group (Fig 5C). This observation was further confirmed through validation with PNGaseF treatment (Fig 5D). Subsequently, utilizing fluorescence microscopy and flow cytometry, we visualized significant enhancements in NLPR3-NEK7 complex formation in 293T cells transfected with vaccine-Env or virulent-Env (Pearson’s coefficient: vaccine-Env: virulent-Env = 0.71:0.96) (Fig 5E). Furthermore, we validated the formation of the NLPR3-NEK7 complex specifically within the Golgi apparatus and the endoplasmic reticulum, rather than in the mitochondrion (Figs 6A, 6B and S3), in 293T cells treated with Nigericin or transfected with either vaccine-Env or virulent-Env. Importantly, we observed that elevated K^+^ concentrations had no effect on the interactions of Env with NLRP3 or NEK7 (S2 Fig). These findings suggest that compared to virulent-Env, vaccine-Env attenuates NLRP3-IL-1β-axis activation by weakening the association between NEK7 and NLRP3.

**Fig 5.**
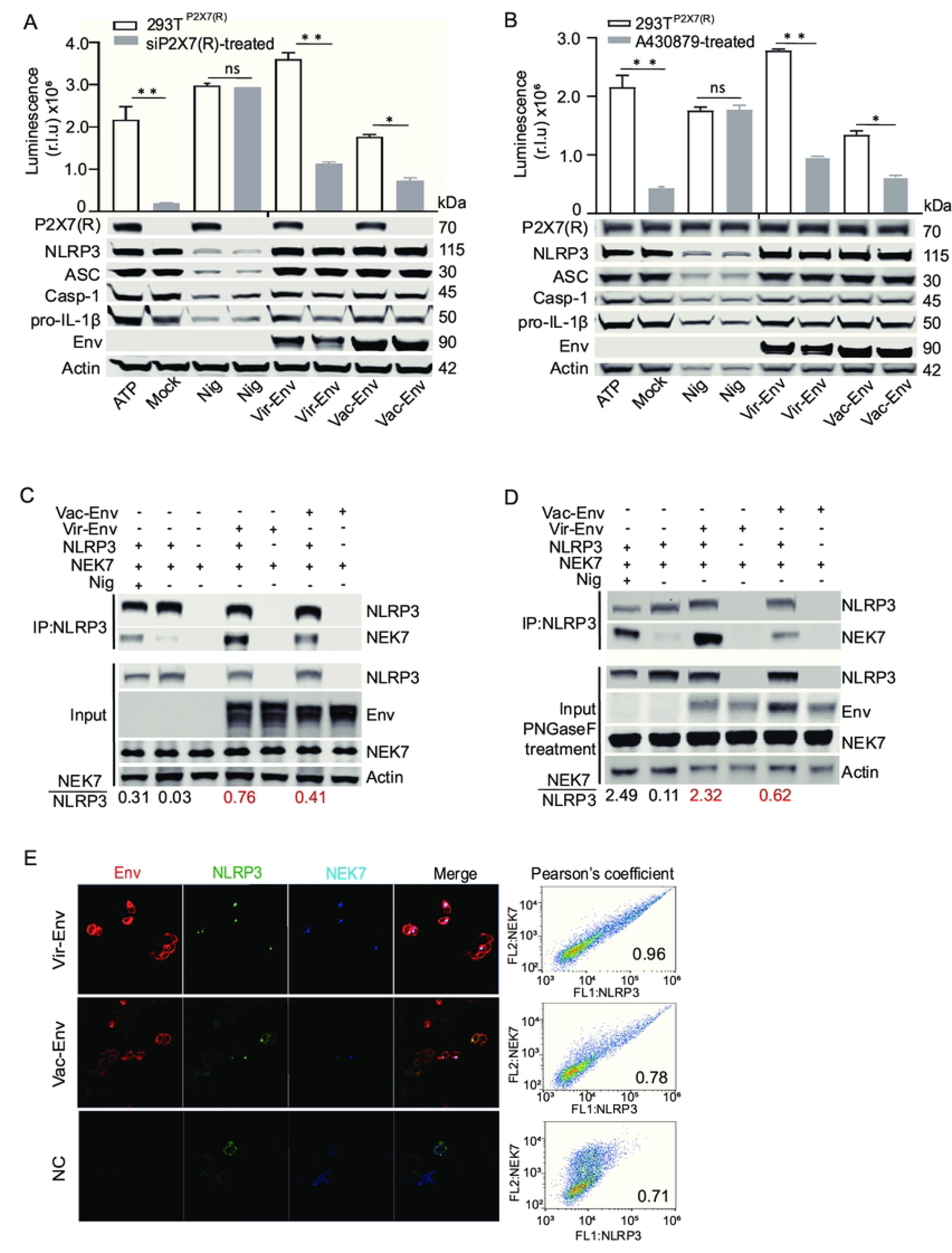
Interaction between NLRP3 and NEK7 in 293T cells transfected with virulent-*env* or with vaccine-*env.* (**A**) Evaluation the activation of the NLRP3-IL-1β axis in 293T^P2X7(R)^ cells transfected with control siRNA or P2X7(R)-specific siRNA and then stimulated with virulent-*env* or vaccine-*env*. (**B**) Evaluation the activation of the NLRP3-IL-1β axis in 293T^P2X7(R)^ cells transfected with virulent-*env* or with vaccine-*env* in the absence or presence of A430879 (a P2X7(R) inhibitor). (**C**) Co-immunoprecipitation of HA-NEK7 or HA-NLRP3 with Flag-EIAV-*env* in 293T cells transfected with the indicated expression plasmids. (**D**) Same as (C) but samples were treated with PNGase F (Glycerol-free) for 1 h before immunoprecipitation. (**E**) The assembly of GFP-NLRP3 and HA-NEK7 in 293T cells co-transfected with virulent-*env* or vaccine-*env*.

**Fig 6.**
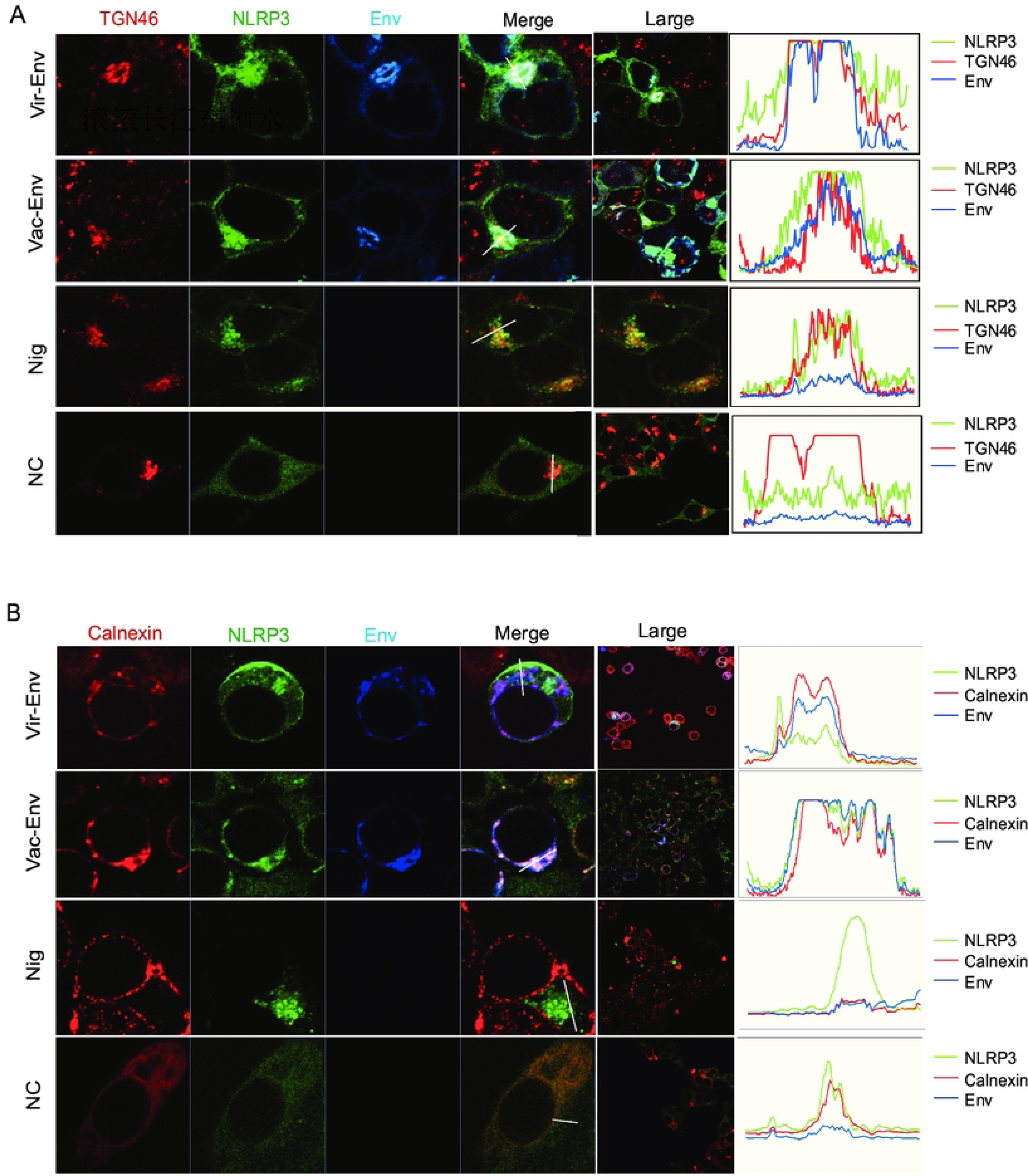
Visualization of NLRP3 inflammasome assembly in 293T cells transfected with EIAV-*env* (s). (**A-B**) Confocal micrograph of assembly of GFP-NLRP3 and HA-NEK7 in TGN46-labeled Golgi apparatus (A) and in Calnexin-labeled endoplasmic reticulum (B) in 293T cells co-transfected with virulent-*env* or vaccine-*env*.

### Four amino acid residues in Env contribute to the differential regulation of the NLRP3-IL-1β-axis by the vaccine and virulent EIAV strains

We aimed to identify the differences between the virulent-Env and the vaccine-Env that were responsible for the apparent differential activation of the NLRP3-IL-1β-axis. To address this, we aligned the amino acid sequences of *env* from virulent and vaccine strains of EIAV to identify any genetic variations. Four key amino acids (S235R, 236D-, N237K and N246K) in Env-Gp90 differed between the vaccine and the virulent EIAV [26, 27] (Fig 7A). We then conducted site-directed mutagenesis in *gp90* and generated four mutants (Fig 7B). Remarkably, the mutant (vir-mut) successfully attenuated the virulent Env (vir-wt), aligning its activity closely with that of the vac-wt. Conversely, this mutant (vir-mut) also enhanced the activity of the attenuated Env (vac-mu) to a level comparable to that of the vir-wt (Fig 7C). These results underscore the functional impact of specific mutations within Env on NLRP3-IL-1β-axis activation, shedding light on the intricate mechanisms underlying the differential immunomodulatory properties of vaccine-Env and virulent-Env. To further validate these findings, we generated libraries pseudotyped with vaccine-Env, virulent-Env, and Env mutants. Encouragingly, consistent results were observed in macrophages infected by these Env-pseudotyped viruses (Fig 7D). We next co-transfected three plasmids (*Env*, *NLPR3* and *NEK7*) into 293T cells, and validated the precipitation of NLPR3-NEK7 in the 293T cells. We found that the interaction between NEK7 and NLRP3 decreased upon vir-mut-Env stimulation, compared with stimulation from vir-wt-Env. Conversely, a stronger interaction between NEK7 and NLRP3 was observed following treatment with vac-mut-Env than with vac-wt-Env (Fig 7E and 7F). These collective findings suggest a decisive role for the four screened amino acid residues in the differential regulation of the NLRP3-IL-1β-axis following infection by the vaccine strain compared to the virulent strain. The underlying mechanisms by which these four amino acid residues contribute to the modulation of the NLRP3-IL-1β-axis will be the focus of future investigations.

**Fig 7.**
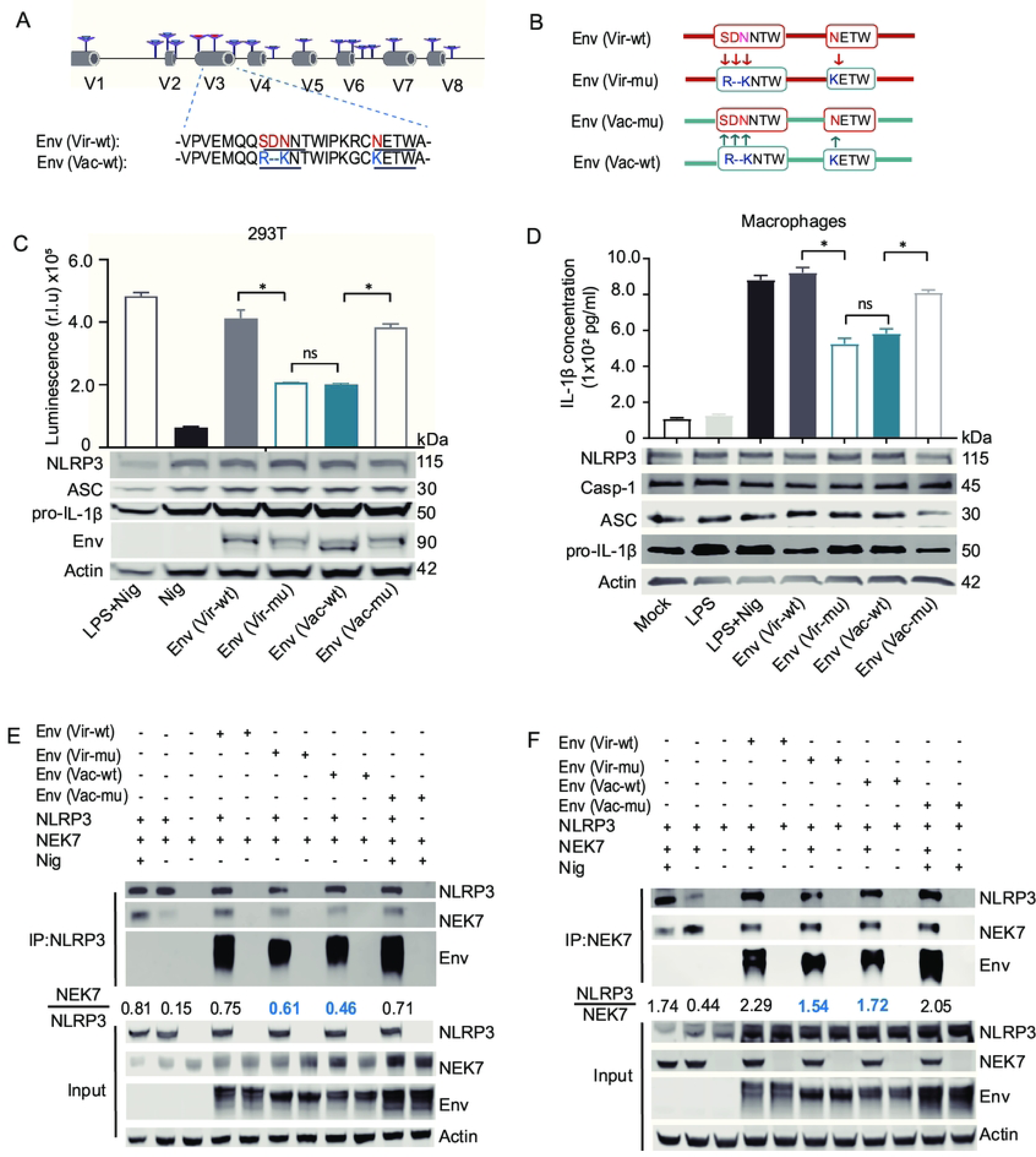
Four amino acid residues are critical for the attenuation of the NLRP3-NEK7 complex assembly induced by vaccine EIAV. (**A**) Schematic representation of the series of mutations constructed by substituting the indicated amino acids, which differ between virulent-*env* (*gp90*) and vaccine-*env* (*gp90*). (**B**) The amino acid sequences of virulent-*env* and vaccine-*env* are written in red and blue, respectively. (**C**) Statistical analyses from Western blot and luciferase activity assays demonstrate the expression levels of IL-1β in 293T cells transfected with EIAV strains (virulent-env, vaccine-env or one of their mutants) using the NLRP3 screening system.(**D**) Representative images and statistical analyses from Western blot and ELISA demonstrate the expression levels of IL-1β in equine macrophages treated with Env-pseudotyped viruses (vaccine-Env, virulent-Env and their mutants) or with LPS and Nigericin as a positive control.(**E-F**) The interactions of NEK7 and NLRP3 with virulent-Env, vaccine-Env or their indicated mutants were analyzed using Co-IP and reciprocal Co-IP assays.

**Fig 8.**
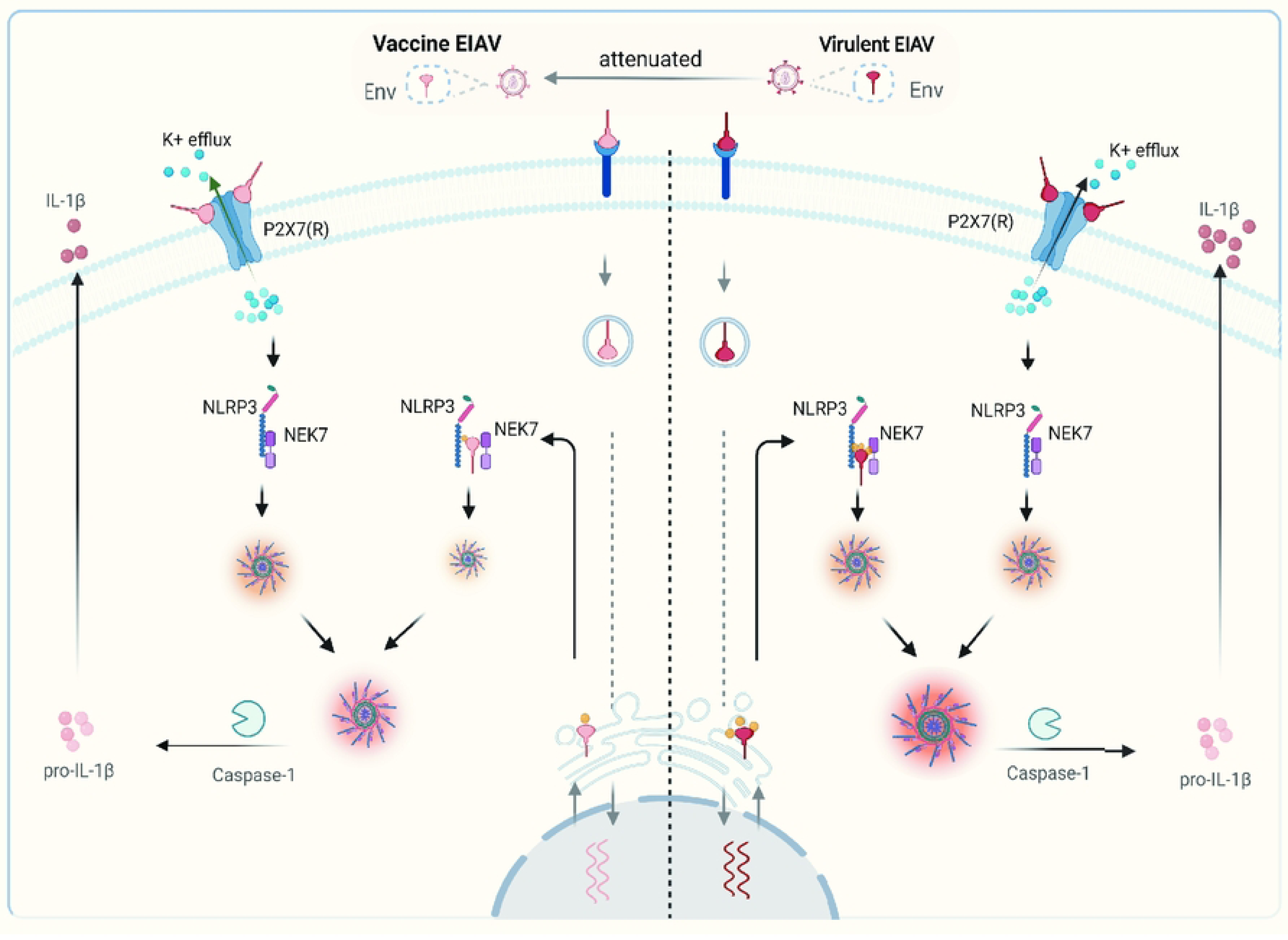
Schematic representation of the mechanism underlying the attenuation NLRP3-IL-1β-axis induced by EIAV vaccine. Both vaccine-Env and virulent-Env mediate NLRP3-IL-1β-axis activation via two-dependent steps. At the attachment stage of EIAV, Env (s) were able to directly bind to P2X7(R) to induce K^+^ efflux. There was no significant difference between two groups at the stage of the Env-P2X7(R) interaction. After the virus enters the cells, expressed virulent-Env and vaccine-Env proteins directly bind to NEK7 and NLRP3 to promote the NEK7-NLRP3 interaction, leading to the assembly and activation of NLRP3 inflammasome. During this stage, vaccine-Env interacted only loosely and sparsely with NLRP3 and NEK7, attenuating NLRP3-IL-1β-axis activation compared to its virulent counterpart.

## Discussion

The findings from your previous studies on horses inoculated with vaccine EIAV compared to those infected with the virulent strain are intriguing. It’s fascinating that the vaccine strain led to reduced IL-1β levels and alleviated inflammatory pathologies [4, 14]. As no assay *in vitro* to examine the NLRP3 inflammasome regulation by EIAV, we initially constructed a platform to systematically visualize the NLRP3-IL-1β axis in cells. The system was able to identify the molecular target of EIAV that responds to the host NLRP3-IL-1β axis. Using this platform, we first identified the molecular target of EIAV-Env that responded to the host NLRP3-IL-1β-axis. Next, we identified K^+^ efflux associated with the Env-P2X7(R) interaction. Interestingly, Env from either the virulent or the attenuated vaccine strain of EIAV was able to interact with P2X7(R), and leading to only non-significant differences in K^+^ ion efflux between the two groups. At the intracellular level, the formation of the NLRP3-NEK7 complex and subsequent pathway activation triggered by Env-NLRP3 and/or Env-NEK7 binding highlight intricate molecular interactions underlying the inflammasome pathway. The proposition that Env-NLRP3 and/or Env-NEK7 binding brings NLRP3 and NEK7 proteins into close proximity, thereby promoting the aggregation of the NLRP3-NEK7 complex in cells, provides valuable insight into the mechanistic aspects of EIAV-induced inflammation. The significant differences observed in the Env-mediated NLRP3-NEK7 complex between the vaccine and virulent EIAV strains, with vaccine-Env interacting weakly and sparsely with NLRP3 and NEK7, further emphasize the potential mechanisms underlying the differential modulation of NLRP3-dependent IL-1β activation by the two strains.

Systematic sequencing of the vaccine and highly virulent EIAV strains allowed us to identify an evolution-dependent adaption in the vaccine strain [8, 13]. Such “natural” evolution-dependent adaption of an EIAV strain could theoretically be as a result of the delicate regulation of activation of the host inflammatory and immune responses, resulting in a decrease in pathological severity. Interestingly, although the amino acid sequences vary between vaccine-Env and virulent-Env, both Env proteins can still bind NLRP3 and NEK7. However, the formation of the NLRP3-NEK7 complex is triggered to different extents by the different viral strains. Therefore, the molecular regulation of NLRP3-NEK7 complex formation by EIAV-Env cannot be explained purely by differences in sequence. To investigate the response of the NLRP3 inflammasome to both vaccine and virulent strains of EIAV, we focused on mutating four amino acids known to differ between the two strains. Mutating the vaccine strain’s Env protein to mimic that of the virulent strain led to increased activation of the NLRP3-IL-1β axis to levels comparable to the virulent strain. Conversely, mutating the virulent strain’s Env to resemble that of the vaccine strain resulted in reduced activation of the NLRP3-IL-1β axis. Hence, we propose that these four amino acids in the vaccine strain of EIAV play a role in finely tuning the NLRP3-IL-1β signaling pathway. Our previous studies predicted that the N246K mutation in wild-type Env results in the loss of an N-glycosylation site in the avirulent vaccine strain. Additionally, deletion of the amino acids at 236 and the N237K mutation also impacts an N-glycosylation site [26, 27]. We therefore propose that the different numbers of N-glycosylation sites between the virulent-Env and vaccine-Env might be responsible for the differential NLRP3 activation and IL-1β expression seen between the two EIAV strains. However, the precise molecular role of these N-glycosylation sites in modulating NLRP3-IL-1β activation warrants further investigation [28].

Our team has demonstrated that the high diversity in the Env of the EIAV vaccine strain leads to greater protective immunity against viral pathogenesis both *in vitro* and *in vivo* [13, 22, 26]. This observation aligns with similar findings reported by other researchers [6, 29, 30], further supporting the stress-dependent flexibility of the immune response against lentiviruses. Therefore, the intricate regulation observed in the vaccine-induced response likely reflects the outcome of a finely tuned evolutionary interplay between the virus and its host [31–33]. As noted by various studies, viruses such as HIV, a well-studied member of the lentivirus family, continuously evolve to establish long-term undetectable latency in response to the stress imposed by efficient HAART therapy [34, 35]. EIAV appears to adopt a more dynamic approach, evolving flexibly rather than lethally to achieve long-term persistence and co-evolution with its equine host.

In conclusion, the EIAV attenuated vaccine presents a representative stable and delicate variation characterized with glycosylation, leading to an accurate and adaptable activation on NLRP3-IL-1β-axis using an Env-P2X7(R)-mediated transcellular signaling as well as Env-NLRP3-NEK7-mediated intracellular signaling. Such special proximity-dependent interaction might benefit a “win-win” adaption during EIAV infection [8]. As no HIV vaccine had been successfully obtained largely due to high glycosylated nature and mutation rate [6, 7, 35, 36], our evolution-based findings might offer an alternative strategy on how HIV vaccine being practically in an unexpected route.

## Materials and methods

### Ethics statement

The horses used in this study were approved by the Harbin Veterinary Research Institute (HVRI), the Chinese Academy of Agricultural Sciences (CAAS). All physical procedures associated with this work were done under anesthesia to minimize pain and distress in accordance with the recommendations of the Ethics Committees of HVRI. The Animal Ethics Committee approval number is Heilongjiang-SYXK (Hei) 2017-009.

### Viruses and Cells

Two EIAV strains were used in our study: i) EIAV_DLV34_ was derived from a cell culture-adapted EIAV virulent strain, which resulted in acute EIA in all the experimentally infected horses at 2 x 10^5^ TCID_50_. ii) The attenuated EIAV vaccine strain (EIAV_DLV121_) was developed from a natural virulent EIAV strain by successive passages in equine monocyte-derived macrophages (eMDMs), and is here termed the vaccine strain. Equine monocyte-derived macrophages and 293T cells were used for infection and transfection separately. Equine monocyte-derived macrophages (eMDMs) were enriched from whole blood as described previously [3, 4]. 293T cells (ATCC, CRL3216) were cultured on DMEM (Sigma-Aldrich, D0819) containing 10 % (vol/vol) fetal bovine serum (FBS; Wisent, 086–15) and antibiotics (1 d% penicillin/streptomycin, Thermo Fisher Scientific, 15,070,063).

### Reagents and antibodies

The following antibodies were used in this study: mouse anti-equine NLRP3 mAb, rabbit anti-equine Caspase-1 pAb, rabbit anti-equine ASC pAb, rabbit anti-equine IL-1β pAb, mouse anti-equine IL-1β mAb and mouse anti-Env mAb were prepared by our laboratory. MCC950 sodium, A430879 and bzATP were purchased from MCE (HY15488, HY-12815 and HY-136254, USA). Glybenclamide (Gly) was purchased from Novus Biologicals (NBP2-30141, USA). Lipopolysaccharide (LPS) (L2280), ATP (A7699), and Mito-TEMPO (SML0737) were purchased from Sigma-Aldrich (St. Louis, MO, USA). TGN46, Tom20 and Calnexin were purchased from Abcam (ab174280, ab186735 and ab133615, USA). Nigericin was obtained from MCE (HY-127019, USA). NEK7 antibody (1:1000) was purchased from Abcam. Antibodies against Flag (F3165) (1:2000) and monoclonal mouse anti-Actin (G9295) (1:5000) were purchased from Sigma (St Louis, USA). Secondary antibodies included DyLight 680 Labeled Anti-Rabbit IgG Antibody (35568, Invitrogen); DyLight 800 Labeled Anti-Rabbit IgG Antibody (KPL,072-06-15-16, Invitrogen); Goat Fluor 488 labeled anti-Rabbit IgG (A-11034, Invitrogen); Alexa Fluor 647 labeled Goat anti-Rabbit IgG (A-2124, Invitrogen); Goat anti-Mouse IgG cross-adsorbed secondary antibody (A-1101, Invitrogen). Lipofectamine 2000 and Lipofectamine™ RNAiMAX transfection reagent were purchased from Invitrogen Corporation (11668019 and 13778075, USA). The equine IL-1β ELISA kit was obtained from the R&D system (DY3340, USA). The IPG-SPEC Assay Kit was purchased from ION Biosciences (3011F, USA). Fluo-4 AM (Calcium Ion Fluorescent Probe, 5mM) was purchased form Beyotime (S1056)

### Transfection and infection

For transfection, 293T cells were seeded into 6-well plates (2 x 10^5^ cells/well) and singly or co-transfected with siRNA oligos (50 nM) using Lipofectamine™ RNAiMAX transfection reagent (Invitrogen) or the indicated plasmids DNA using Lipofectamine 2000 (Invitrogen) according to the manufacturer’s instructions. The expression of these target proteins was confirmed using western blotting 24 hours after transfection. For virus infection, equine macrophages were enriched from whole blood and were cultured as previously reported [3]. The eMDMs were cultured for one day and then infected with EIAV strains at 2 x 10^5^ TCID_50_. Cells and supernatants were collected at the indicated time-points. The expression of the inflammasome components was confirmed using western blotting. Supernatants were collected for analysis of mature IL-1β using an ELISA kit (R&B, USA).

### Histopathology and Pathological assessment

For the histopathology studies, three horses were inoculated with attenuated vaccine and three horses were infected with virulent strain as previously described [3, 22]. Thirty days after inoculation, the horses were euthanized and the organs were collected and examined. Liver, lung, spleen, kidney and lymph node of each horse were embedded in paraffin for 24 h, then sliced and stained with haematoxylin and eosin (H&E). Semi-quantitative analysis of inflammatory pathological changes in each organ was scored independently as previously described [22]. Pathological assessments of five autopsied organs were performed by three independent professional pathologists in a double-blind experiment.

### Assay for NLRP3-IL-1β-axis activation and IL-1β measurement

The NLRP3-IL-1β-axis screening system, which monitors the cleavage of pro-IL-1β, was developed to study equine NLRP3 inflammasome activation based on the technique published in a previous report [37]. This reporter is based on the biological activity of a pro-IL-1β-*Gaussia* luciferase (iGluc) fusion. Briefly, the pro-IL-1β-dependent formation of a protein complex renders the iGluc enzyme inactive. The cleavage of pro-IL-1β leads to monomerization of this biosensor protein, and the fusion protein then gains strong luciferase activity. According to the working principle of this system, expression plasmids coding for eqcaspase-1, eqASC, eqNLRP3 and eqIL-1β were constructed using the INFUSION cloning technique (Clontech). 293T cells (2 x 10^5^ cells/well) were then transiently transfected with the inflammasome plasmid constructs with a total of 250 ng DNA per 12-well plate. At 24 h post-transfection, the supernatant of the cultured cells was measured to assess the luciferase signal using Coelenterazine (Sigma, 55779-48-1). Adherent cells were lysed in passive lysis buffer (E1941, Promega) so that the western blot assay could be run in parallel. IL-1β measurement was carried out with an equine IL-1β ELISA kit (R&B, USA).

### Real-time RT-PCR

Total RNA was extracted using TRizol (Invitrogen) according to the supplier’s instructions. Real-time RT-PCR was performed using the SYBR-Green method as described previously using available primers. cDNA was amplified for the IL-1β gene (sense: 5’-GAGGAGGATGGCCCAAAACA-3’; anti-sense: 5’-AGC CACAATGATTGACACGAC-3’), the NLRP3 gene (sense: 5’-TCGGTTGGTGAACTG CTCTC-3’; anti-sense: 5’-CGGTGAGGCTCCAGTTAGTG-3’), the NLRC4 gene (sense: 5’-AAGTCTCTGTCAGCCGAACC-3’; anti-sense: 5’-CGAGACTGCTCTCCTTCAGT-3’), and the NLRP1 gene (sense: 5’-CACACCTGGAAAGGAATCAGAGA-3’; anti-sense: 5’-TCCGGGGTCAGATGTGTGTA-3’). Actin mRNA (sense: 5’-CATCTGCTG GAAGGTGGACAA-3’; anti-sense: 5’-CGACATCCGTAAGGACCGTTA-3’) was examined as the cellular reference control. The relative expression of each gene was calculated using delta-delta -Ct method (2-ΔΔCt).

### Western blots and immunoprecipitation

At 24 h after transfection as described above, cells were lysed in an appropriate cell lysis buffer (1% Triton X-100, 50 mM Tris-HCl, pH 7.4, and 150 mM NaCl) with 1 % Protease Inhibitor Cocktail (HY-K0010, MCE). After running the lysates on a 4 %-12 % or 12 % SDS-PAGE gel, proteins were transferred onto a PVDF membrane. The blots were probed with primary antibodies at RT for 1-2 hours and then incubated with the indicated secondary antibodies (1:10000). For immunoprecipitations, cell pellets were lysed and incubated overnight with the appropriate antibodies plus anti-Flag or anti-HA magnetic beads (Sigma-Aldrich, A2095) as described previously. Signals were detected using a Licor Odyssey 448 imaging system (USA). For relative protein expression, images were quantified in ImageJ software and normalized using the corresponding endogenous β-actin expression.

### Confocal Microscopy

For immunostaining, 293T cells or primary eMDMs were seeded on coverslips and transfected (or infected) with the indicated plasmids (or EIAV strain). After 24 h post-transfection, or following infection at 6 h, 12 h and 24 h, cells were fixed in 4 % paraformaldehyde (P0099, Biotime) for 15 min and subjected to saponin permeabilization (P0050, Biotime). Cells were then incubated with the corresponding antibodies and suitable secondary antibodies. Nuclei were stained with DAPI (P0131, Beyotime) for 10 min. All cells were washed with PBS 5 times after each step. Fluorescence images were acquired using a confocal laser scanning microscope (Carl Zeiss LSM 800), and analyzed using ZEN 2 (blue edition) software.

### Mass spectrometry

The eMDMs were separated and cultured as described previously [4], and then were infected with an EIAV strain. Three days post-infection, cells were collected by scraping into a lysis buffer (1% Triton X-100, 50 mM Tris-HCl, pH 7.4, and 150 mM NaCl). The cell lysates were cleared using centrifugation (15,000 r.p.m.,10 min). Dilution buffer and then normal mouse IgG2b and anti-Env magnetic beads (MCE, HY-K0202) were added into the supernatant, and incubated for 2 h at 4 °C with rotation. The supernatant was removed, and the beads were washed extensively with 1 x PBST. An epitope peptide targeting Env was then added (1mg/ml) to disrupt the interaction between Env and the beads. The eluted samples were loaded on a large 12 % SDS-PAGE (BioRad, 4561068) and resolved, and were then analyzed using western blotting. Electrophoresis was run for longer (40-50 min) than usual to ensure good band resolution. The stained protein bands were then cut out and incubated in deionized water for further mass spectrometric analysis. Mass spectrometry was performed by the BGI Genomics company. Briefly, each sample was dehydrated, reduced, alkylated, and then digested with trypsin. The peptides were extracted and delivered onto a nano RP column and eluted with a gradient. Peptide identification was carried out in the Mascot software (Version 2.3.01, Matrix Science, UK) using the UniProt database search algorithm and the integrated false discovery rate (FDR) analysis function. The data were searched against a protein sequence database downloaded from 2019 uni_horse horse (50,340 sequences; 30,790,782 residues).

### EIAV-Env pseudovirus construction

Briefly, the packaging constructs psPAX2 (AIDS Resource and Reagent Program), luciferase reporter plasmid Plenti-CMV Puro-Luc (Addgene), and the plasmids expressing the virulent-Env, vaccine-Env or Env mutants were used for EIAV-Env pseudovirus generation. These plasmids were co-transfected into 293T cells (5 x 10^4^ cells/well) using calcium phosphate. After 48 h transfection, supernatants were collected and infectivity was determined with reverse transcriptase activity assays (Sigma, 11468120910).

### RNAi

293T cells were transfected with a pool of 3 P2X7 (R) -targeting siRNAs (TGACAGAAATTGACAACAA; AGACAAGAACAACTCCAAA; ACAGTGTCTTAACATTCAA) or control siRNA (CAAACAGAAUGGUCCUAAA) (50 μM) using Lipofectamine™ RNAiMAX transfection reagent (Invitrogen, 13778100) according to the manufacturer’s instructions. The efficiency of siRNA silencing of P2X7(R) was evaluated using western blotting.

### Generation of the stable P2X7(R) 293T cell line

Briefly, the pSIN4-P2X7 lentivirus was generated by co-transfecting 293T cells with packaging plasmids psPAX2 and pMD2.G. Medium containing viruses was filtered and added to the target cells. 24 h following initial infection with the lentiviruses, cells were selected on puromycin (1µg/µl) (Gibco, A1113803) for at least 14 days. The remaining cells were enriched using fluorescence-activated cell sorting (FACS) and were then expanded until there were enough cells for validation with immunoblotting.

### Measurement of intracellular potassium

Intracellular K^+^ concentrations were determined with a potassium probe (IPG-2AM) according to the manufacturer’s instructions. Briefly, equine macrophages were cultured and plated in a 96-well microplate (1.5 x 10^5^ cells/well) as described previously [4]. On the day of the experiment, equine macrophages were first infected with either EIAV virulent or vaccine strains separately at the same TCID_50_, and were then either treated with Nigericin (10 µM) (Sigma, 28643-80-3) at the indicated time-points (0 min, 15 min, 30 min, 45 min, 60 min, 120 min, 240 min and 360 min), or left untreated. 100 µl/well potassium probe was then added. After 1 h incubation at 37 °C, fluorescent signals were detected using an ELISA plate reader, with excitation/emission wavelengths of 520/540 nM. All reagents in this assay were provided in the IPG-SPEC Assay Kit (3011F, ION Biosciences).

### Measurement of intracellular Ca^2+^

Intracellular Ca^2+^ concentrations were measured in equine macrophages using the Ca^2+^ sensitive dye, Fluo-4-AM. Briefly, equine macrophages were infected with either EIAV virulent or vaccine strains at the same dose. At the indicated timepoints, cells were switched and washed three times with wash buffer and incubated with 10 μM Fluo-4-AM for 45 min at 37 °C, followed by three washes and an additional incubation in PBS for 20 min. The fluorescence value (excitation at 488 nm) was monitored using a laser scanner, and the emitted light signal was read at 510 nm.

### Flow cytometry

293T cells (1 x 10^6^) were co-transfected with plasmids expressing NLRP3, NEK7 and virulent-Env or vaccine-Env separately. Cells were then collected and stained with a 1:200 dilution of each of the following antibodies: Alexa Fluor 647 anti-rabbit IgG (Invitrogen, A-21244) and Fluor 405 anti-mouse IgG (Invitrogen, A-31556). Samples were fixed with BD Cyto fixation buffer (BD Biosciences, 554655) before sorting on the BD LSRFortessa (BD). Analysis was performed using the FlowJo software version 10.

### Statistical analysis

Image J software was used to measure the band intensity from the western blot. GraphPad Prism was used for statistical tests. All results are expressed as mean ± SD. Treatment groups were compared using two-tailed, unpaired Student’s t-tests (Prism, GraphPad). Error bars represent SD (standard error of the mean). ns, not significant (*P* > 0.05), \**P* < 0.05, \*\**P* < 0.01, \*\*\**P*< 0.001.

## Acknowledgments

We would like to thank Professor Xin Yin for helpful discussions. We thank Peng Zhang for assistance on animal work, and the Core Facility of the Harbin Veterinary Research Institute, the Chinese Academy of Agricultural Sciences for providing the technology platform. This study was supported by grants from Heilongjiang Provincial Natural Science Foundation of China (LH2023C049 to Y L), the National Natural Science Foundation of China (32372985 to Y L), the Tianchi Talent Introduction Plan and the National Key Research and Development Program of China (2023YFD1802500).

## Author contributions

X.J.W. conceived and supervised the study and revised the manuscript; Y.Z.L conceptualized the project and wrote the paper. X.G, C.L and H.X.L designed and supervised the experiments. Y.H.W performed the evaluation of pathological results. H.L.R and L.N helped construct mutants and generated Env-pseudotyped viruses. S.W.M supplied 293T^NEK7^ stable cells. All authors discussed the results and commented on the manuscript.

## Data and software availability

All data are available upon request to the lead contact. No proprietary software was used in the paper.

## Figure Legends

**S1 Fig.** Comparison of inflammatory pathological changes in specimens from horses inoculated with the EIAV vaccine and horses infected with the virulent EIAV strain. Typical inflammatory pathological changes observed in lung, kidney, liver, spleen and lymph gland on infection with EIAV vaccine or virulent strains are presented separately (haematoxilin and eosin 4 m paraffin sections, original magnification 10x). Severity of pathological lesions and scoring are given in Table 1.

**S2 Fig.** K+ efflux affects the interaction between NEK7 and NLRP3, but not that between Env and NLRP3 or NEK7. (**A**) 293T cells were co-transfected with NEK7-Flag and NLRP3-HA in the presence of 50 μM concentrations of KCl (increasing intracellular K+ concentration). Immunoprecipitation and analyses were as described in Fig 3D. (**B-C**) Procedure was as in (A) but instead of transfection with NEK7 and NLRP3, cells were co-transfected with *env*-HA and NLRP3-Flag (B) or co-transfected with *env*-HA and NEK7-Flag (C) at high concentrations of KCl (50 μM). An immunoprecipitation assay was then performed using the indicated antibodies.

**S3 Fig.** Visualization of NLRP3 inflammasome localization in 293T cells transfected with EIAV-*env* (s). Confocal micrograph of assembly of GFP-NLRP3 and HA-NEK7 not Tom20-labeled mitochondria in 293T cells co-transfected with virulent-*env* or vaccine-*env*.

## Reference

1. Haas L. [Equine infectious anemia--a review]. Berl Munch Tierarztl Wochenschr. 2014;127(7-8):297–300. (Die equine infektiose Anamie--eine Ubersicht.) PMID: 25080822.

2. Koumangoye R. The role of Cl(-) and K(+) efflux in NLRP3 inflammasome and innate immune response activation. Am J Physiol Cell Physiol. 2022;322(4):C645–C52. Epub 20220216. 10.1152/ajpcell.00421.2021 PMID: 35171697.

3. Lin YZ, Shen RX, Zhu ZY, Deng XL, Cao XZ, Wang XF, et al. An attenuated EIAV vaccine strain induces significantly different immune responses from its pathogenic parental strain although with similar in vivo replication pattern. Antiviral Res. 2011;92(2):292–304. Epub 20110825. 10.1016/j.antiviral.2011.08.016 PMID: 21893100.

4. Lin YZ, Cao XZ, Li L, Li L, Jiang CG, Wang XF, et al. The pathogenic and vaccine strains of equine infectious anemia virus differentially induce cytokine and chemokine expression and apoptosis in macrophages. Virus Res. 2011;160(1-2):274–82. Epub 20110714. 10.1016/j.virusres.2011.06.028 PMID: 21782860.

5. Cohn LB, Chomont N, Deeks SG. The Biology of the HIV-1 Latent Reservoir and Implications for Cure Strategies. Cell Host Microbe. 2020;27(4):519–30. 10.1016/j.chom.2020.03.014 PMID: 32272077; PubMed Central PMCID: PMCPMC7219958.

6. Rolland M. HIV-1 phylogenetics and vaccines. Curr Opin HIV AIDS. 2019;14(3):227–32. 10.1097/COH.0000000000000545 PMID: 30925535.

7. Chun TW, Fauci AS. Latent reservoirs of HIV: obstacles to the eradication of virus. Proc Natl Acad Sci U S A. 1999;96(20):10958–61. 10.1073/pnas.96.20.10958 PMID: 10500107; PubMed Central PMCID: PMCPMC34225.

8. Wang X, Wang S, Lin Y, Jiang C, Ma J, Zhao L, et al. Genomic comparison between attenuated Chinese equine infectious anemia virus vaccine strains and their parental virulent strains. Arch Virol. 2011;156(2):353–7. Epub 20101207. 10.1007/s00705-010-0877-8 PMID: 21136127.

9. Zimmerli U, Thur B. [Equine Infectious Anaemia - a review from an official veterinary perspective]. Schweiz Arch Tierheilkd. 2019;161(11):725–38. 10.17236/sat00232 (Die Equine Infektiose Anamie - eine Besprechung aus amtstierarztlicher Sicht.) PMID: 31685446.

10. Lohmann KL, James CR, Higgins SN, Howden KJ, Epp T. Disease investigations for equine infectious anemia in Canada (2009-2012) - Retrospective evaluation and risk factor analysis. Can Vet J. 2019;60(11):1199–206. PMID: 31692681; PubMed Central PMCID: PMCPMC6805026.

11. Wang HN, Rao D, Fu XQ, Hu MM, Dong JG. Equine infectious anemia virus in China. Oncotarget. 2018;9(1):1356–64. Epub 20170821. 10.18632/oncotarget.20381 PMID: 29416700; PubMed Central PMCID: PMCPMC5787444.

12. Cursino AE, Vilela APP, Franco-Luiz APM, de Oliveira JG, Nogueira MF, Junior JPA, et al. Equine infectious anemia virus in naturally infected horses from the Brazilian Pantanal. Arch Virol. 2018;163(9):2385–94. Epub 20180511. 10.1007/s00705-018-3877-8 PMID: 29752558.

13. Wang XF, Lin YZ, Li Q, Liu Q, Zhao WW, Du C, et al. Genetic Evolution during the development of an attenuated EIAV vaccine. Retrovirology. 2016;13:9. Epub 20160203. 10.1186/s12977-016-0240-6 PMID: 26842878; PubMed Central PMCID: PMCPMC4738788.

14. Liu Q, Ma J, Wang XF, Xiao F, Li LJ, Zhang JE, et al. Infection with equine infectious anemia virus vaccine strain EIAVDLV121 causes no visible histopathological lesions in target organs in association with restricted viral replication and unique cytokine response. Vet Immunol Immunopathol. 2016;170:30–40. Epub 20160123. 10.1016/j.vetimm.2016.01.006 PMID: 26832985; PubMed Central PMCID: PMCPMC7112881.

15. Groslambert M, Py BF. Spotlight on the NLRP3 inflammasome pathway. J Inflamm Res. 2018;11:359–74. Epub 20180925. 10.2147/JIR.S141220 PMID: 30288079; PubMed Central PMCID: PMCPMC6161739.

16. Xu H, Akinyemi IA, Chitre SA, Loeb JC, Lednicky JA, McIntosh MT, et al. SARS-CoV-2 viroporin encoded by ORF3a triggers the NLRP3 inflammatory pathway. Virology. 2022;568:13–22. Epub 20220117. 10.1016/j.virol.2022.01.003 PMID: 35066302; PubMed Central PMCID: PMCPMC8762580.

17. Rossi C, Salvati A, Distaso M, Campani D, Raggi F, Biancalana E, et al. The P2X7R-NLRP3 and AIM2 Inflammasome Platforms Mark the Complexity/Severity of Viral or Metabolic Liver Damage. Int J Mol Sci. 2022;23(13). Epub 20220704. 10.3390/ijms23137447 PMID: 35806450; PubMed Central PMCID: PMCPMC9267345.

18. Wang W, Hu D, Feng Y, Wu C, Song Y, Liu W, et al. Paxillin mediates ATP-induced activation of P2X7 receptor and NLRP3 inflammasome. BMC Biol. 2020;18(1):182. Epub 20201126. 10.1186/s12915-020-00918-w PMID: 33243234; PubMed Central PMCID: PMCPMC7694937.

19. Rosli S, Kirby FJ, Lawlor KE, Rainczuk K, Drummond GR, Mansell A, et al. Repurposing drugs targeting the P2X7 receptor to limit hyperinflammation and disease during influenza virus infection. Br J Pharmacol. 2019;176(19):3834–44. Epub 20190819. 10.1111/bph.14787 PMID: 31271646; PubMed Central PMCID: PMCPMC6780046.

20. Sikora A, Liu J, Brosnan C, Buell G, Chessel I, Bloom BR. Cutting edge: purinergic signaling regulates radical-mediated bacterial killing mechanisms in macrophages through a P2X7-independent mechanism. J Immunol. 1999;163(2):558–61. PMID: 10395640.

21. Ward JR, West PW, Ariaans MP, Parker LC, Francis SE, Crossman DC, et al. Temporal interleukin-1beta secretion from primary human peripheral blood monocytes by P2X7-independent and P2X7-dependent mechanisms. J Biol Chem. 2010;285(30):23147–58. Epub 20100521. 10.1074/jbc.M109.072793 PMID: 20495003; PubMed Central PMCID: PMCPMC2906308.

22. Lin Y, Wang XF, Wang Y, Du C, Ren H, Liu C, et al. Env diversity-dependent protection of the attenuated equine infectious anaemia virus vaccine. Emerg Microbes Infect. 2020;9(1):1309–20. 10.1080/22221751.2020.1773323 PMID: 32525460; PubMed Central PMCID: PMCPMC7473056.

23. Ma J, Wang SS, Lin YZ, Liu HF, Wei HM, Du C, et al. An attenuated EIAV strain and its molecular clone strain differentially induce the expression of Toll-like receptors and type-I interferons in equine monocyte-derived macrophages. Vet Microbiol. 2013;166(1-2):263–9. Epub 20130621. 10.1016/j.vetmic.2013.06.005 PMID: 23850441.

24. Adinolfi E, Giuliani AL, De Marchi E, Pegoraro A, Orioli E, Di Virgilio F. The P2X7 receptor: A main player in inflammation. Biochem Pharmacol. 2018;151:234–44. Epub 20171227. 10.1016/j.bcp.2017.12.021 PMID: 29288626.

25. Pacheco PAF, Faria RX. The potential involvement of P2X7 receptor in COVID-19 pathogenesis: A new therapeutic target? Scand J Immunol. 2021;93(2):e12960. Epub 20200908. 10.1111/sji.12960 PMID: 32797724; PubMed Central PMCID: PMCPMC7461012.

26. Liu C, Wang XF, Wang Y, Chen J, Zhong Z, Lin Y, et al. Characterization of EIAV env Quasispecies during Long-Term Passage In Vitro: Gradual Loss of Pathogenicity. Viruses. 2019;11(4). Epub 20190424. 10.3390/v11040380 PMID: 31022927; PubMed Central PMCID: PMCPMC6520696.

27. Han X, Zou J, Wang X, Guo W, Huo G, Shen R, et al. Amino acid mutations in the env gp90 protein that modify N-linked glycosylation of the Chinese EIAV vaccine strain enhance resistance to neutralizing antibodies. Viral Immunol. 2010;23(5):531–9. 10.1089/vim.2009.0006 PMID: 20883167.

28. Son S, Hwang I, Han SH, Shin JS, Shin OS, Yu JW. Advanced glycation end products impair NLRP3 inflammasome-mediated innate immune responses in macrophages. J Biol Chem. 2017;292(50):20437–48. Epub 20171019. 10.1074/jbc.M117.806307 PMID: 29051224; PubMed Central PMCID: PMCPMC5733583.

29. Goudsmit J, Back NK, Nara PL. Genomic diversity and antigenic variation of HIV-1: links between pathogenesis, epidemiology and vaccine development. FASEB J. 1991;5(10):2427–36. 10.1096/fasebj.5.10.2065891 PMID: 2065891.

30. Korber B, Hraber P, Wagh K, Hahn BH. Polyvalent vaccine approaches to combat HIV-1 diversity. Immunol Rev. 2017;275(1):230–44. 10.1111/imr.12516 PMID: 28133800; PubMed Central PMCID: PMCPMC5362114.

31. Sorin M, Kalpana GV. Dynamics of virus-host interplay in HIV-1 replication. Curr HIV Res. 2006;4(2):117–30. 10.2174/157016206776055048 PMID: 16611052.

32. Ahlers JD. All eyes on the next generation of HIV vaccines: strategies for inducing a broadly neutralizing antibody response. Discov Med. 2014;17(94):187–99. PMID: 24759623.

33. Dhamija N, Rawat P, Mitra D. Epigenetic regulation of HIV-1 persistence and evolving strategies for virus eradication. Subcell Biochem. 2013;61:479–505. 10.1007/978-94-007-4525-4_21 PMID: 23150264.

34. Kulkosky J, Bray S. HAART-persistent HIV-1 latent reservoirs: their origin, mechanisms of stability and potential strategies for eradication. Curr HIV Res. 2006;4(2):199–208. 10.2174/157016206776055084 PMID: 16611058.

35. Castro-Gonzalez S, Colomer-Lluch M, Serra-Moreno R. Barriers for HIV Cure: The Latent Reservoir. AIDS Res Hum Retroviruses. 2018;34(9):739–59. Epub 20180828. 10.1089/AID.2018.0118 PMID: 30056745; PubMed Central PMCID: PMCPMC6152859.

36. Rashid A, Li K, Feng Y, Ahmad T, Getaneh Y, Yu Y, et al. HIV-1 genetic diversity a challenge for AIDS vaccine development: a retrospective bibliometric analysis. Hum Vaccin Immunother. 2022;18(1):2014733. Epub 20220111. 10.1080/21645515.2021.2014733 PMID: 35016590; PubMed Central PMCID: PMCPMC8973384.

37. Bartok E, Bauernfeind F, Khaminets MG, Jakobs C, Monks B, Fitzgerald KA, et al. iGLuc: a luciferase-based inflammasome and protease activity reporter. Nat Methods. 2013;10(2):147–54. Epub 20130106. 10.1038/nmeth.2327 PMID: 23291722; PubMed Central PMCID: PMCPMC4127477.

